# Effect of heme oxygenase-1 on the expression of interferon-stimulated genes

**DOI:** 10.1101/2024.11.04.620611

**Authors:** Patryk Chudy, Katarzyna Bednarczyk, Eryk Chatian, Wojciech Krzeptowski, Agata Szade, Krzysztof Szade, Monika Żukowska, Jan Wolnik, Grzegorz Sokołowski, Alicja Józkowicz, Witold N. Nowak

## Abstract

Heme oxygenase-1 (HO1, *Hmox1*) degrades excess heme and is considered an anti-oxidative and anti-inflammatory enzyme. Our previous studies in *Hmox1* knockout mice revealed induction of interferon-stimulated genes (ISGs) in all cell types analyzed, despite unchanged interferon production. Here, we sought to identify the pathway underlying HO1-dependent ISG regulation and determine how ISG expression changes in cultured cells in response to stressors typical of *Hmox1*-deficient mice.

Using murine wild-type and *Hmox1*-deficient (KO-Hmox1) fibroblasts, we showed that in cells cultured under control conditions, the expression of most of the tested ISGs was independent of cellular HO1 status. We then analyzed the effect of extrinsic stressors: hemolytic, oxidative, genotoxic, and replication stress, proinflammatory TNFα, and endogenous heme overload. TNFα (upregulated in *Hmox1* knockout mice) was the sole and universal ISG inducer in both wild-type and KO-Hmox1 fibroblasts. Unexpectedly, the response of KO-Hmox1 cells to exogenous TNFα was weakened, probably due to impaired NF-κB activity and reduced p65 nuclear retention. A similar decrease we observed for STAT1. Additionally, the presence of TREX1 exonuclease in the nucleus indicated impaired nuclear envelope integrity. Noteworthy, HO1 colocalizes with PARP1, a protein regulating cytoplasmic-nuclear transport. Olaparib-mediated PARP1 inhibition abolished TNAα-induced nuclear accumulation of p65 and STAT1 in wild-type cells, but not in KO-Hmox1 counterparts.

In summary, the inflammation typical of *Hmox1*-deficient mice appears to be a major inducer of ISGs *in vivo*. Despite this, the inflammatory response to exogenous TNFα is attenuated in KO-Hmox1 cells, likely due to decreased nuclear retention of NF-κB and STAT1.

## Introduction

Heme oxygenase-1 (HO1, encoded by *Hmox1* gene) is an enzyme that degrades heme and affects many cellular processes, including DNA replication, cell cycle, and cell differentiation [1], [2], [3]. In contrast to constitutive HO2 (encoded by *Hmox2* gene), it is an inducible form, upregulated under stressful conditions. HO1 is known for its cytoprotective, antioxidant, anti-inflammatory, and proangiogenic properties, which are exerted either by heme degradation products (biliverdin, Fe^2+^ ions, and carbon monoxide) [4], or regulation of heme availability [3], [5], or by direct HO1 protein binding to transcription factors [6]. Modulation of HO1 expression or activity has, however, very different effects depending on the cellular context [7].

Our team has shown that HO1 in the bone marrow niche is necessary to protect hematopoietic stem cells (HSCs) from premature aging [8]. By comparing the transcriptome of wild-type and *Hmox1^-^*^/-^ mouse primary HSCs, bone marrow-derived endothelial cells (ECs), and CXCL12-abundant reticular cells (CARs) [8], as well as muscle satellite cells (SCs) [9], we found that there is no single pattern of gene expression changes characteristic for *Hmox1* deficiency. However, one of the groups of genes with similarly altered expression in all cell types examined are interferon-I (IFN-I) stimulated genes (ISGs) (Fig. 1A). This relationship is indeed prominent: top 4 upregulated genes in *Hmox1*^-/-^ HSCs are *Ifi44, Ifi204*, and *Ifi27*, while among top 4 upregulated genes in *Hmox1*^-/-^ CARs are *Oas1g, Oas1a*, and *Ifi27l2a* [8]. Generally, in *Hmox1^-/-^* mice, we observed increased expression of interferon-regulatory factors (IRFs), interferon-induced proteins (IFIs), interferon-induced proteins with tetratricopeptide repeats (IFITs), and 2’-5’-oligoadenylate synthetases (OASs) genes. Yet, genes coding for interferons and their receptors were unaffected [8] (Fig. 1B).

**Figure 1.**
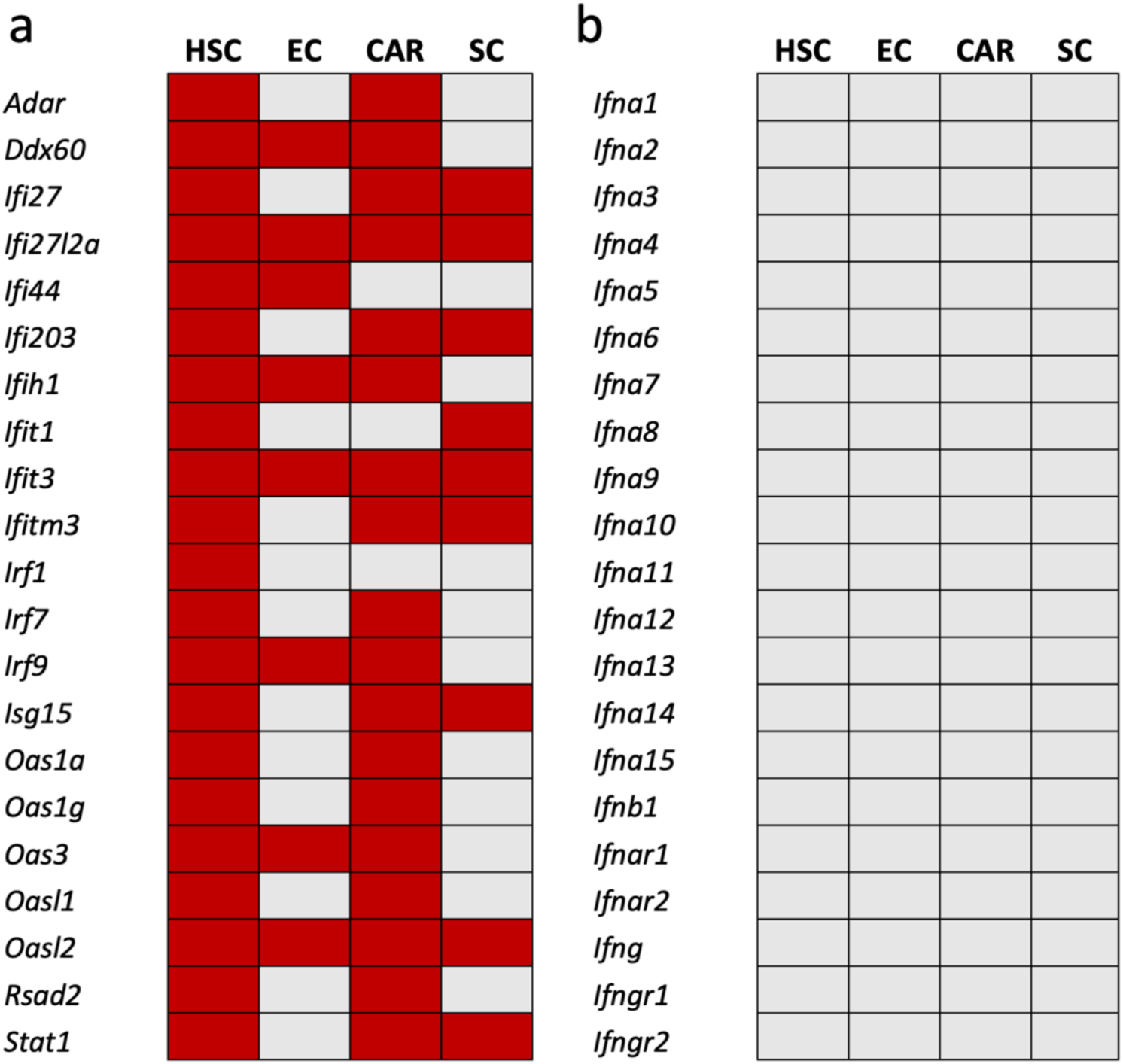
Schematic representation of RNA-seq analysis. Differential expression of **a)** several IFN-I regulated genes and **b)** genes encoding interferons and their receptors. Red – up-regulation in *Hmox1^-/-^* cells, gray – no differences. HSC – hematopoeitic stem cells; EC – endothelial cells; CAR – CXCL12-abundant reticular cells; SC – muscle satelite cells.

Type I interferons are family of cytokines representing a part of antiviral defense systems, that includes IFN-β and a group of very similar IFN-α proteins [10], [11]. They share a common type I interferon receptor (IFNAR), a heterodimer consisting of IFNAR1 and IFNAR2 subunits. IFN-α is produced in particularly high levels by mononuclear phagocytes, whereas IFN-β rather is expressed by nonhematopoietic cells [12]. The major stimulus for IFN-I expression is a viral infection. The initial recognition of virus-derived nucleic acids and the first wave of IFN-I production are mainly mediated by pattern recognition receptors (PRRs), such as endosomal Toll-like receptor (TLR) 7 and 9 [13]. Then, the detection of viral replication products is mediated by cytosolic receptors such as DDX58 or DDX60 helicase [14].

Upon TLR7 and TLR9 activation, signal is transduced through IRFs, mainly through IRF3 and IRF7, which then translocate from the cytoplasm to the nucleus to promote the transcription of IFN-α and IFN-β [15], [16]. TLR7 and TLR9 receptors may also activate NF-κB transcription factor to trigger inflammatory response [17]. Type I interferons bind to IFNAR, thereby activating the receptor associated TYK2 and JAK1 tyrosine kinases. This results in phosphorylation of STAT1 and STAT2 proteins. STATs bind to IRF9 and form a heterotrimeric transcription factor ISGF3, that recognizes IFN-stimulated response elements (ISRE) in target genes [18]. A group of these genes, together with IRF7 and IRF9, master regulators of IFN-I dependent cellular responses, are strongly upregulated in different *Hmox1*-deficient primary cells [8], [9]. One can expect that such an effect is regulated at the cell-autonomous level, as HO1 protein has been shown to directly interact with IRF3, influencing its activity [19], [20]. However, this has not been verified so far.

Additionally, our recent work on the protective role of HO1 in replication stress showed that HO1 affects the expression and function of PARP1 [3] – an enzyme that transfers ADP-ribose moieties from nicotinamide adenine dinucleotide (NAD^+^) to a plethora of proteins in response to DNA damage, cellular stress, and inflammation, in a process called PARylation [21]. PARP1 and other members of PARP family modulate IFN-I response [22]. PARylation also enhances the activity of NF-κB [23] and its retention in the nucleus [22]. On the other hand, mono-ADP-ribosylation (MARylation) by PARP10, another member of the family, negatively regulates NF-κB signaling [24]. Finally, PARylation is crucial for the activity of STAT1, which forms both ISFG [25] and homodimers activating IRF1 and IRF8 [26]. Therefore, we suspected that the cellular effects of HO1 deficiency on IFN-I response genes might be associated with the HO1-PARP1 axis.

It should be emphasized, however, that comparisons of gene expression profiles of primary cells isolated directly from wild-type and *Hmox1* knockout (KO) mice [5], [8] do not allow for distinguishing between cell-autonomous effects and responses to external signals originating from the *Hmox1*-deficient microenvironment or the whole organism. Both in mice and humans, lack of HO-1 is associated with susceptibility to oxidative stress, enhanced DNA damage response, and development of progressive inflammatory disease, which all may affect ISGs [8], [19], [27], [28]. Therefore, distinguishing intrinsic from extrinsic effects on gene expression profile would require analyzing of ISGs in cells cultured *in vitro* in homeostatic conditions, and then examining their response to extrinsic stressors.

We aimed to elucidate whether dysregulation of interferon-stimulated genes also occurs in *Hmox1*-deficient cells cultured *in vitro* and to determine how the regulation of ISG expression changes in response to stressors typical of *Hmox1* deficiency.

## Materials and Methods

### Animals

C57BL/6J×FVB Hmox1^KO^ (KO-Hmox1) and C57BL/6J×FVB Hmox1^WT^ (WT) mice were bred in the animal facility of the Faculty (Breeder Registry nr 078). We used them only for tissue collection, and therefore there was no ethical committee approval required. Mice were housed in individually ventilated cages in specific pathogen-free conditions and had unlimited access to food and water. Mice were euthanized via CO_2_ inhalation.

### Primary fibroblast isolation and cell culture

For the isolation of fibroblasts, we used KO-Hmox1 and WT mice bred in our animal facility. Mice were euthanized via CO_2_ inhalation. Fibroblasts were isolated from mouse tails by digestion in collagenase II (2.5 mg/mL in DPBS, Gibco) for 90 minutes and then filtrated by using 70 μm cell strainer (Biologix). Cell cultures were performed under standard conditions, at 37°C in a humidified atmosphere with 5% CO_2_. Fibroblasts were cultured in DMEM High-Glucose medium (Biowest) containing 10% fetal bovine serum (FBS, EurX), antibiotics (100 IU/mL penicillin and 100 µg/mL streptomycin, Sigma-Aldrich) and fibroblast growth factor 2 (FGF2; 10 ng/mL, PeproTech). Cells were cultured for a maximum of 3 passages in T-25 flasks (Falcon) and then seeded into 6-well or 24-well plates (Falcon) on glass coverslips.

In experiments, cells were treated with 10 ng/mL TNFα (a kind gift from Dr. Krystyna Stalińska, Department of Cell Biotechnology, FBBB, Jagiellonian University), 10 ng/mL aphidicolin (APH, A0781-1MG, Sigma-Aldrich), 0.25 µM etoposide (ETO, E1383-250MG, Sigma-Aldrich), 350 µM 5-aminolevulinic acid (ALA, A7793-500MG, Sigma-Aldrich), 20 mg/mL hemoglobin (Hb, H2500, Sigma-Aldrich), 100 µM H_2_O_2_ (95321-500ML, Sigma-Aldrich), 10 µM NF-κB pathway inhibitor (INH14, MedChem Express), 10 µM AP-1 pathway inhibitor (T5224, MedChem Express) or 0.1 µM olaparib (HY-10162, MedChem Express).

### Immunofluorescence staining

Cells grown on glass coverslips covered with 1% Geltrex LDEV-Free Reduced Growth Factor Basement Membrane Matrix (Gibco) were fixed with 4% Pierce methanol-free formaldehyde (Thermo Scientific) at room temperature for 10 minutes, followed by two washes with PBS (Lonza). Cells fixed in formaldehyde were permeabilized in 0.2% Triton X-100 in PBS (PBS-Tx) at room temperature for 10 minutes and blocked in 10% normal donkey serum (Sigma-Aldrich) in 0.1% PBS-Tx at room temperature for 30 minutes. Then, the cells were incubated overnight at 4°C with primary antibodies diluted 1:200 in 0.1% PBS-Tx with 1% donkey serum. Staining was followed by three PBS washes before incubation with the appropriate secondary antibodies that were diluted 1:400 in 0.1% PBS-Tx containing 1% donkey serum at room temperature for one hour. Finally, after washing with PBS and counterstaining the cell nuclei with 0.5 µg/mL DAPI (Sigma-Aldrich) for 10 minutes at room temperature, the coverslips were mounted in Fluorescence Mounting Medium (Dako) and allowed to dry before imaging. Negative controls with the omission of primary antibodies were performed for each protein. Cells were analyzed using the Axio Observer Z1 microscope (Zeiss) with 63x/1.4 Oil DIC m27 objective or using Leica DMI6000 with 40x/0.75 objective.

### Antibodies

The following antibodies were used: anti-HO-1 (ADI-SPA-894, Enzo), anti-TNFR1 (BS-2941R, Thermo Scientific), anti-phospho-IκB-α (Ser32/36) (9246, Cell Signaling), anti-NF-κB p65 (51-0500, Thermo Scientific), anti-phospho-NF-κB p65 (Ser536) (MA5-15160, Thermo Scientific), anti-STAT1 (PA5-95442, Thermo Scientific), anti-STAT2 (MA5-42463, Thermo Scientific), anti-lamin A/C (4C11, Abcam), anti-exportin-1/CRM1 (D6V7N) (46249T, Cell Signaling Technology), anti-tubulin hFAB rhodamine (12004166, Bio-Rad), anti-rabbit StarBright Blue 700 (12004161, Bio-Rad), anti-rabbit HRP linked antibody (7074P2, Cell Signaling Technology), anti-rabbit Alexa Fluor 488 (A21206, Thermo Scientific).

### Analysis of NR4A1 in mouse hematopoietic stem cells (HSCs)

Bone marrow was isolated from femurs and tibias of 6-month-old male individuals as described earlier [8]. Bone marrow was filtered through a cell strainer (100 µm) and centrifuged at 670 *g* for 10 minutes at 4°C. Next, the cell pellet was resuspended in an RBC lysis buffer (0.15 mol/L NH4Cl, 10 mmol/L KHCO3, 0.1 mmol/L EDTA) and incubated for 7 minutes at room temperature. After diluting the samples with 2% FBS in PBS, the cells were centrifuged. Finally, the cell pellet was resuspended with 100 µL of 2% FBS in PBS. Mouse HSCs (Lin^-^c-Kit^+^Sca-1^+^ (LKS) CD150^+^ CD48^-^), MPP (LKS CD150^-^ CD48^-^) and GMP (LKS CD150^-^CD48^+^) were stained using following clones of antibodies: anti-mouse CD3, clone 17A2; anti-mouse Ly-6G/Ly-6C, clone RB6-8C5; anti-mouse CD11b, clone M1/70; anti-mouse CD45R/B220, clone RA3-6B2; anti-mouse TER-119/erythroid cells, clone Ter-119; anti-mouse-CD150, clone TC15-12F12.2; anti-mouse-CD48, clone HM-48-1 (all Biolegend); anti-mouse-Ly6A/E (Sca-1), clone D7; anti-mouse-CD117 (c-Kit), clone 2B8 (eBioscience). Then, cells were fixed and permeabilized with BD IntraSure kit (BD Biosciences) and stained with anti-Nurr77(Nr4a1) antibody (clone JM59-11, Thermo Scientific), and then donkey anti-rabbit AlexaFluor®568 antibody (Thermo Scientific). Signal for NR4A1 antibody was collected in PE-Texas Red channel on BD LSR Fortessa flow cytometer. Secondary only control (omitting primary Nurr77/NR4A1 antibody) was used to set the gating strategy.

### Detection of oxidative stress

CellROX Deep Red Reagent (Thermo Fisher Scientific) was used for the general oxidative stress detection. Cells were cultured in 12-well plates. Tert-butyl hydroperoxide (TBHP, 200 µM) was added for 30 minutes as a positive control. To detect reactive oxygen species, CellROX Deep Red reagent (500 nM) was added to each sample and incubated for 30 minutes at 37°C. Cell fluorescence was analyzed using LSR Fortessa flow cytometer (Becton Dickinson).

### Lipid Peroxidation

Click-iT Lipid Peroxidation (LAA) Kit for Imaging – Alexa Fluor 488 (C10446, Thermo Fisher Scientific) was used, following the provider’s protocol. Cells were cultured in 24-well plates with round coverslips and stimulated with 50 µM linoleamide alkyde (LAA) for 24 h. Cumene hydroperoxide (CH) served as a positive control. Then, the cells were fixed with 4% Pierce methanol-free formaldehyde (Thermo Scientific) at room temperature for 10 minutes and washed twice with PBS. The cells were permeabilized in 0.1 % PBS-Tx at room temperature for 10 minutes. Lipid peroxidation was detected by incubation of coverslips with 50 µL Click-iT reaction cocktail for 30 minutes at room temperature. Then, the samples were washed 3 times with PBS and co-stained with DAPI (0.5 µg/mL; Sigma-Aldrich). Fluorescence detection was performed using Leica DM6B fluorescence microscope with PL Fluotar L 20x/0.40 objective.

### Reverse Transcription and Real-Time PCR

RNA was isolated using a RNeasy Mini Kit (Qiagen) and reverse transcribed with a QuantiTect Reverse Transcription Kit (Qiagen) with integrated gDNA removal. The gene expression was assessed on a StepOnePlus thermocycler (Applied Biosystems) with real-time PCR using an SYBR Green JumpStart Taq ReadyMix (Sigma-Aldrich) and specific primers: *Eef2* (F: GCG GTC AGC ACA ATG GCA TA, R: GAC ATC ACC AAG GGT GTG CAG), *Irf1* (F: GGA TAT GGA AAG GGA CAT AAC, R: ATA AGG TCT TCG GCT ATC TTC), *Irf7* (F: TAA GGT GTA CGA ACT TAG CC, R: TAC TGC AGA ACC TGT GTG), *Irf9* (F: CTA CTT CTG TAG AGA TTT GGC, R: GAT GAG ATT CTC TTG GCT ATG), *Ifitm3* (F: AAC TTC TGA GAA ACC GAA AC, R: ATC TCA GCC ACC TCA TAT TC), *Adar1* (F: CAT CAG GTT TCT CTA CAG TG, R: CTG CAG GAT TTG TCA AAG AG), *Oas1g* (F: TCA ATG TCG TGT GTG ATT TC, R: CTG GTG AGA TTG TTA AGG AAC), *Oasl1* (F: CTC CTC TGT ATC TAC TGG AC, R: CCA CTA TGT CCC ATC TGT AG).

### Immunoblotting

Cultured cells were detached with TrypLE (Gibco), suspended in cold PBS (Lonza) and centrifuged at 400 *g* for 10 minutes. Then, the pellets were resuspended in Pierce RIPA buffer (Thermo Scientific) with protease inhibitors (complete Protease Inhibitor Cocktail, Merck) and incubated for 5 minutes at 4°C with agitation. Lysates were clarified by centrifugation at 8000 *g* for 10 minutes at 4°C. Protein concentration was determined by a BCA assay kit (Thermofisher) to ensure the equal loading of each sample (10 μg of proteins). Samples were electrophoretically separated in 4–20% Mini-PROTEAN TGX Precast Protein Gels (BioRad) followed by transfer to a nitrocellulose blotting membrane in transfer buffer with 20% ethanol (Trans-Blot Turbo RTA Mini 0.2 µm Nitrocellulose Transfer Kit, BioRad). The membranes were blocked with EveryBlot Blocking Buffer (BioRad) for 5 minutes and incubated with primary antibodies overnight at 4°C. The membranes were then washed three times with TBST and incubated with secondary anti-rabbit-HRP or anti-rabbit StarBright Blue 700 and anti-tubulin hFAB rhodamine antibodies diluted in EveryBlot blocking buffer for 1 hour at room temperature. After five washes with TBST, the membrane was optionally incubated with horse-radish peroxidase (HRP) substrate (Bio-Rad) (if an HRP-linked antibody was used). Immunofluorescence or chemiluminescence detection was performed on a ChemiDoc MP instrument (Bio-Rad).

### Trans-AM ELISA

Cells were seeded on 6-well plates (100 000 cells/well) and treated with 10 ng/mL TNFα for 30 minutes. Then, the cells were detached with TrypLE (Gibco), suspended in cold PBS (Lonza), and centrifuged at 400 *g* for 10 minutes. Nuclear fraction was isolated from cell pellets using Cell Fractionation Kit - Standard (ab109719, Abcam). Then, trans-AM ELISA was done using Transam NF-κB p65 (40096, Active Motif). First, the binding of NF-κB to its consensus sequence was performed by adding 2 μg of each nuclear fraction for 1 h at room temperature on wells activated with Complete Binding Buffer. Then, after washing three times, the anti-NF-κB p65 antibody diluted 1:1000 in Antibody Binding Buffer was incubated for 1 h at room temperature. The reaction was followed with three PBS washes before an incubation with anti-rabbit-HRP secondary antibodies that were diluted 1:1000 in Antibody Binding Buffer and incubated for 1 h at room temperature. Finally, after washing four times, a colorimetric reaction was performed by incubation with Developing Solution for 5 minutes and then with Stop Solution at room temperature. Absorbance was measured within 5 minutes at 450 nm with an optional reference wavelength of 655 nm on a TECAN infinite M200 microplate reader (TECAN).

### Luciferase activity assay

Cells seeded on a 24-well plate (50 000 cells/well) were transfected with 200 ng/well of pNL3.2.NF-κB-RE [NlucP/NF-κB-RE/Hygro] Vector (N1111, Promega) using Lipofectamine 2000 Transfection Reagent (11668027, Thermo Scientific) and incubated for 24 h at 37°C. Then, the cells were treated with 100 ng/mL TNFα for 5 h, detached with TrypLE, centrifuged at 400 *g* for 10 minutes, and resuspended in culture medium. Nano-Glo Luciferase Assay (N1110, Promega) was performed by adding an equal volume of Nano-Glo reagent into cells suspension on black 96-well plate (Perkin Elmer). After 3 minutes of incubation at room temperature, the luminescence signal was measured on a TECAN infinite M200 microplate reader (TECAN).

### Proximity ligation assay (PLA)

For the *in-situ* visualization of the interaction between p65 and CRM1 proteins, we followed the Duolink PLA Fluorescence Protocol using the Duolink In Situ Detection Reagents Orange kit (Sigma-Aldrich). Briefly, the cells grown on glass coverslips covered with 1% Geltrex LDEV-Free Reduced Growth Factor Basement Membrane Matrix (Gibco), were fixed with 4% Pierce methanol-free formaldehyde (Thermo Scientific) and permeabilized with 0.2% PBS-Tx (PBS with 0.2% Triton X-100) for 10 minutes at room temperature. Next, cells were blocked in a drop of Blocking Solution (Sigma-Aldrich) at 37°C for one hour, and incubated with primary antibodies [anti-p65 (sc-8008, Santa Cruz), anti-CRM1 (46249T, Cell Signaling Technology)] diluted 1:200 in Duolink Antibody Diluent (Sigma-Aldrich). After washing in wash buffer A (10 mM Tris, 150 mM NaCl and 0.05% Tween 20), cells were incubated with secondary antibodies. In Situ PLA secondary antibodies (Sigma-Aldrich) were used: anti-mouse PLUS, and anti-rabbit MINUS. Next, the cells were again washed twice in wash buffer A, and then incubated with the ligase (diluted 1:40 in ligation buffer) for 30 minutes at 37°C. After the next round of washing in wash buffer A, cells were incubated with polymerase (diluted 1:80 in an amplification buffer) for 100 minutes at 37°C. Finally, cells were washed in wash buffer B (200 mmol/L Tris and 100 mmol/L NaCl), counterstained with DAPI (0.5 μg/mL, Sigma-Aldrich), mounted in Fluorescence Mounting Medium (Dako), and allowed to dry before imaging. Negative controls were performed using primary and secondary antibodies only. Fluorescence detection was performed using Axio Observer Z1 microscope (Zeiss) using 63x/1.4 Oil DIC m27 objective.

### RNA Seq data

We analyzed our previously published data available at in the BioProject database, accession no. PRJNA562450 [8].

### Cytokine concentration analysis

INFγ, IL-1β, IL-10, MCP1 and TNFα protein concertation in WT and KO-Hmox1 mice blood serum was measured by using Luminex (MILLIPLEX MAP Mouse Cytokine/Chemokine Premixed 32 Plex, Mouse Cytokine/Chemokine Magnetic Bead Premixed 32 Plex and the custom assay panel, Millipore). Assays were performed according to manufacturer’s instructions. The samples were diluted 1:1 in Assay Buffer and incubated with Premixed Beads overnight at 4°C. Signal detection was done using FLEXMAP 3D system (Millipore).

### ELISA

INFα/β protein concertation in mice WT and KO-Hmox1 blood serum was measured using ELISA kit (R&D Systems) according to manufacturer’s instructions. Absorbance was measured on a TECAN infinite M200 microplate reader (TECAN).

### Statistical Analysis

All experiments were performed in duplicate or triplicate and were repeated independently at least three times unless otherwise indicated. Data were analyzed with GraphPad Prism 8.0 software. Tests used in the statistical analysis are listed in the description of the figures. Bar graphs represent mean ± SEM. ns – nonsignificant, *-p≤0.05, **-p<0.01, ***-p<0.001.

## Results

### HO1 deficiency leads to upregulation of *Ifi27* both *in vivo* and *in vitro*

In earlier experiments [8], [9] we observed an increase in mRNA expression of many ISGs but not interferons and their receptors (Fig. 1). Gene expression was, however, examined in cells that are not typical interferon producers, namely HSCs, ECs, and CARs from the bone marrow, and satellite cells from the muscle. We also measured interferon concentration in the blood serum: the levels of INFα/β and INFγ were very low, close to the detection limits, and did not differ significantly between genotypes (Fig. 2A). This confirms that the production of interferons in *Hmox1*-deficient mice was not enhanced, despite the upregulated expression of ISGs. In the same animals, we found a trend towards increased concentration of interleukin-10 (IL-10) and significant elevation of monocyte chemoattractant protein-1 (MCP1), the cytokines induced by IFNα and IFNγ, respectively [29]. This may also indicate activation of ISGs. Additionally, we observed a tendency to increase the concentration of IL-1β and higher level of tumor necrosis factor-α (TNFα), as expected for the proinflammatory state of KO-Hmox1 mice (Fig. 2B).

**Figure 2.**
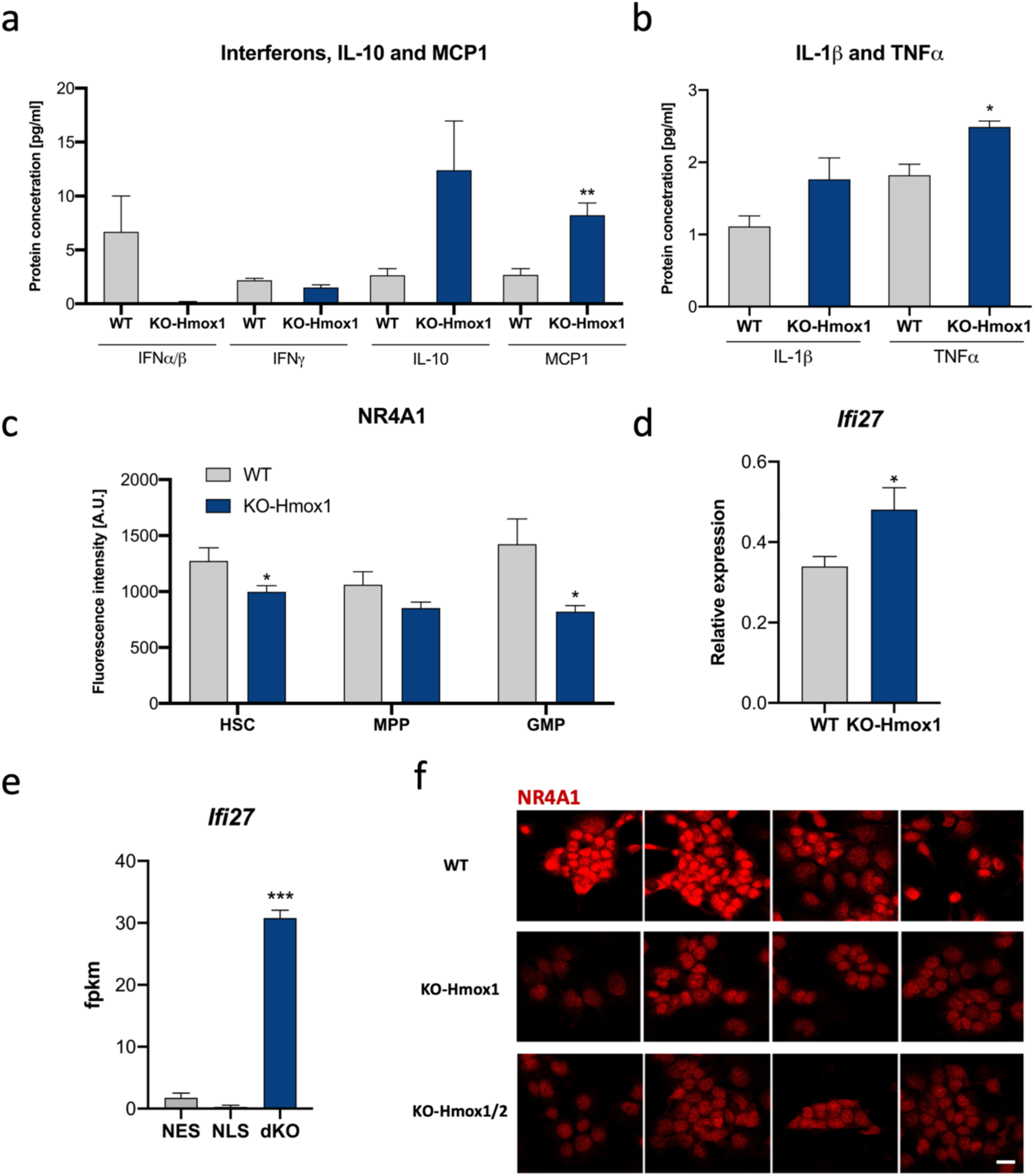
The levels of **(a)** INFα, INFɣ, IL-10 and MCP1 (N=3-6. T-test) and **(b)** IL-1β and TNFα in WT and KO-Hmox1 mice blood serum. N=3-6. T-test. **c)** Flow cytometry analysis of NR4A1 protein levels in WT and KO-Hmox1 HSC, MPP and GMP cells. N=5. T-test. **d)** qPCR. Expression of *Ifi27* gene in WT and KO-Hmox1 fibroblasts under control conditions. N=4. T-test. **e)** RNA-seq. Expression of *Ifi27* gene in dKO, NES, NLS iPSC under control conditions. N=4. T-test **f)** Representative images of NR4A1 immunofluorescence staining in WT, KO-Hmox1 and KO-Hmox1/2 iPSC. Scale bar: 20 µm.

Importantly, it seems that changes in ISG expression have a functional significance. The gene whose expression was universally increased in almost all KO-Hmox1 cells was *Ifi27* and its paralog *Ifi27l2a* (Fig. 1A). IFI27 protein localizes to the inner membrane of the nuclear envelope. It is responsible, among others, for the export of NR4A1 (NURR77) nuclear receptor from the nucleus to the cytoplasm, in an exportin-1 (CRM1)-dependent manner. Export to the cytoplasm leads to proteasomal degradation of NR4A1 and inhibition of NR4A1-dependent pathways. Such regulation is especially important in HSCs because NR4A1 is an essential factor regulating HSC quiescence and metabolism [30]. It can be assumed that increased expression of *Ifi27* would be associated with lower levels of NR4A1 protein. Indeed, we observed a decrease in NR4A1 protein in HSCs and granulocyte-monocyte progenitors (GMPs), with a similar trend in multipotent progenitors (MPPs) isolated from *Hmox1*-deficient mice (Fig. 2C).

To test whether *Ifi27* is also upregulated in other cell types, we used fibroblasts isolated from KO-Hmox1 mice cultured *in vitro* for several passages. RT-PCR analysis showed a slight but significant increase in the expression of *Ifi27* in fibroblasts lacking HO1 (Fig. 2D). In addition, we used induced pluripotent stem cells (iPSCs) that had been subjected to deletion of the endogenous *Hmox1* and *Hmox2* genes and then engineered to express only cytoplasmic or nuclear form of HO1 [3]. Here, we also observed strong upregulation of *Ifi27* in the absence of HO1 (Fig. 2E). Similarly to the results from hematopoietic cells (Fig. 2C), also in iPSCs, the absence of HO1 or both HO1 and HO2 was associated with a decrease in NR4A1 protein levels, confirming the functional significance of the changes in IFI27 expression (Fig. 2F). The increased expression of *Ifi27* not only in *Hmox1*-deficient cells directly isolated from mice but also in those cultured *in vitro* may suggest the existence of intrinsic regulation.

### In cells grown under homeostatic conditions, the expression of many ISGs is independent of HO1 status

All subsequent experiments were performed using *in vitro* cultured fibroblasts, which have relatively high levels of HO1 protein under control conditions (Fig. 3A). In contrast to wild-type (WT) cells, in fibroblasts lacking *Hmox1* (KO-Hmox1), HO1 protein was undetectable (Fig. 3A). Of note, KO-Hmox1 cells did not differ from wild type counterparts in terms of total reactive oxygen species (ROS) levels (Fig. S1A) and lipid peroxidation (Fig. S1B). This means that the lack of HO1 does not cause increased oxidative stress in our experimental setting.

**Figure 3.**
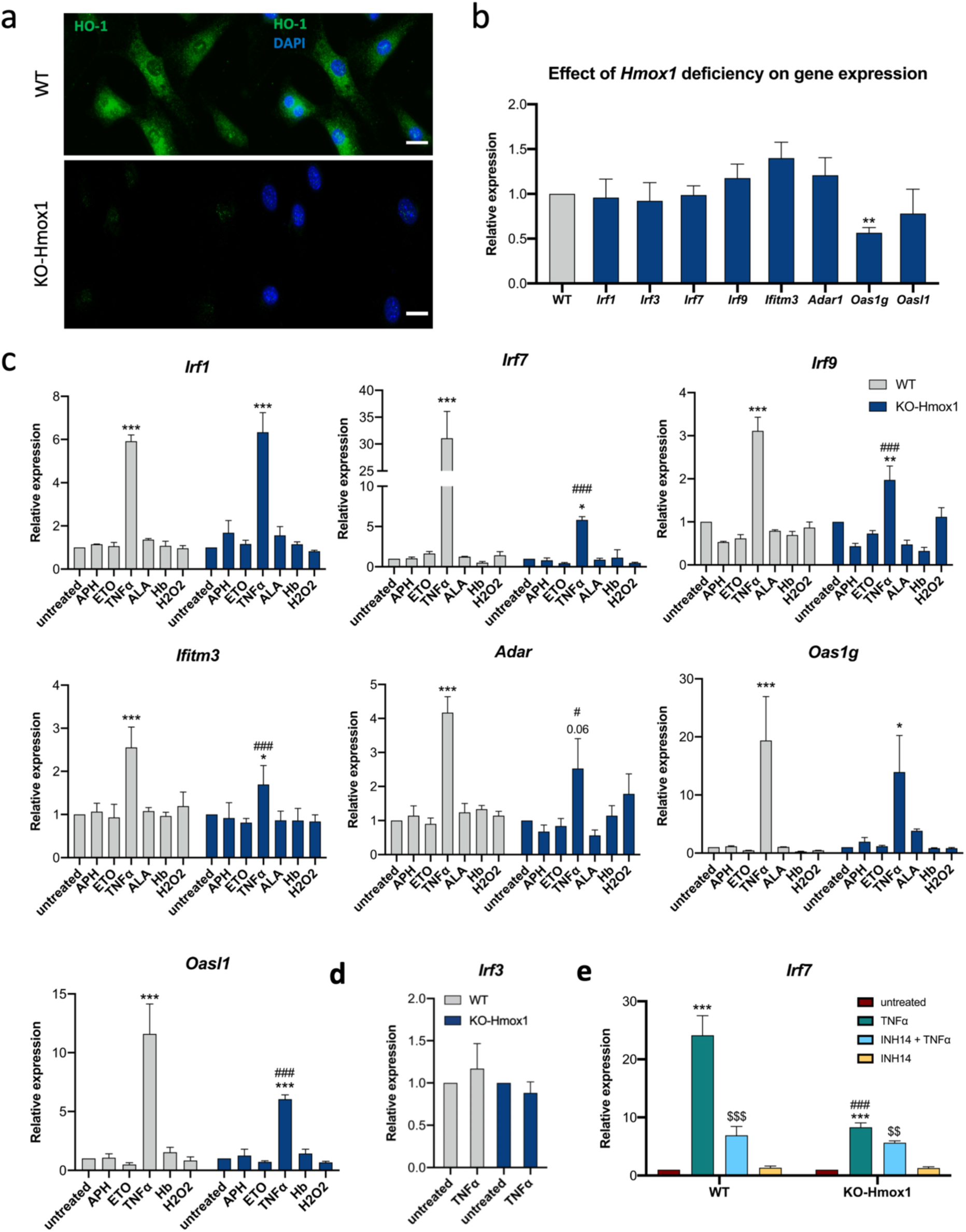
Type-I interferon response genes expression panel (qPCR). **a)** Representative images of HO-1 immunofluorescence staining (green) and nuclei counterstained with DAPI (blue) in WT and KO-Hmox1 fibroblasts. Scale bar: 20 µm. **b)** Relative expression of INF-I genes in KO-Hmox1 fibroblasts (blue) comparing to WT fibroblasts (gray) under control conditions. N=4. One-way ANOVA. **c)** Relative expression of *Irf1, Irf7, Irf9, Ifitm3, Adar1, Oas1g, Oasl1* in WT and KO-Hmox1 fibroblasts, untreated or treated with aphidicolin (APH, 0.01 µg/ml), etoposide (ETO, 0.25 µM), TNFα (10 ng/ml), 5-aminolevulinic acid (ALA, 350 µM), hemoglobin (Hb, 20 mg/ml) and H_2_O_2_ (100 µM) for 24 h. N=3-4. Two-way ANOVA. * - untreated vs treated, # - WT vs KO-Hmox1. Gene expression in treated cells was normalized to control conditions for each genotype separately. **d)** Relative expression of *Irf3* in WT and KO-Hmox1 fibroblasts treated with TNFα (10 ng/ml) for 24 h. N=6. Two-way ANOVA. **e)** Relative expression of *Irf7* in WT and KO-Hmox1 fibroblasts untreated or treated with TNFα (10 ng/ml) and INH14 (inhibitor of NF-κB pathway, 10 µM) both individually or in combination for 24h. N=3. Two-way ANOVA. * - untreated vs TNFα, $ - untreated vs INH14 + TNFα, # - WT vs KO-Hmox1.

First, we examined the influence of genotype on the basal expression of several ISGs, namely *Irf1*, *Irf7*, *Irf9*, *Ifitm3*, *Adar*, *Oas1g* and *Oasl1*. All these genes were upregulated in HSCs and some other cell types isolated from KO-Hmox1 mice (Fig. 1). Under control conditions, unlike *Ifi27* (Fig. 2D), none of the tested genes showed increased expression in *Hmox1*-deficient cells (Fig. 3B). This indicates that the increased expression of ISGs *in vivo* was due to the action of some external stressor(s).

### Expression of ISGs is upregulated by TNFα

In the next step, we stimulated fibroblasts with compounds inducing different types of stress that have been described in *Hmox1*-deficient cells, namely with aphidicolin (APH – replication stress), etoposide (ETO – genotoxic stress), TNFα (inflammatory response), 5-aminolevulinic acid (ALA – enhanced heme synthesis and overload), hemoglobin (Hb – hemolytic stress), and H_2_O_2_ (oxidative stress) for 24 hours. Out of all tested compounds, only TNFα increased the expression of *Irf1, Irf7, Irf9, Ifitm3, Adar, Oas1g* and *OasL1*. Moreover, TNFα was a universal stressor that induced the expression of all tested ISGs in both WT and KO-Hmox1 fibroblasts *in vitro* (Fig. 3C). This indicates that the induction of ISGs observed in various primary cells *in vivo* is mainly extrinsically regulated and suggests that it may be a consequence of the pre-inflammatory state typical of KO-Hmox1 mice, particularly the increased production of TNFα. It is worth noting that the expression of Irf3, which is not regulated at the transcriptional level [31] and which was not induced *in vivo* (data not shown), did not change *in vitro* in response to TNFα (Fig. 3D).

### Response to TNFα is reduced in KO-Hmox1 cells

The regulation of two genes, *Irf1* and *Oas1g*, by TNFα was comparable regardless of HO1 status. Surprisingly, induction of *Irf7, Irf9, Adar, Oas1g* and *OasL1* was significantly lower in the absence of HO1 (Fig. 3C). This suggests that unlike under control conditions (proinflammatory in *Hmox1*-deficient mice), if inflammation is induced, the response to TNFα may be stronger in the presence of HO1.

Induction of ISGs may be regulated by NF-κB [32], [33], which directly targets the expression of *Irf7* [34]. On the other hand, it has been suggested that NF-κB activity may be controlled by HO1, both through its products and via a direct protein-protein interaction between HO1 and the p65 subunit [35]. Therefore, in an attempt to identify the mechanism of the attenuated response to TNFα, we first checked whether NF-κB was indeed involved in the induction of ISGs in our cells.

To this aim we treated WT and KO-Hmox1 fibroblasts with TNFα in the presence or absence of INH14, a specific inhibitor of NF-κB pathway, and then analyzed the expression of *Irf7* (Fig. 3E). INH14 inhibited the response to TNFα stimulation, confirming the involvement of the NF-κB-dependent pathway. However, the effect of the inhibitor was much stronger in WT cells, where activation was reduced by 71% than in KO-Hmox1 cells, where activation was reduced by 32%. Thus, the stimulation of *Irf7* expression in response to TNFα was much stronger in WT cells, but in the presence of the NF-κB inhibitor, it was similar in both genotypes (respectively: p<0.001 and p>0.99, two-way ANOVA). This may indicate that the stronger response to TNFα in the presence of HO1 results from a more efficient action of NF-κB and that this pathway is somehow defective in the *Hmox1*-deficient cells.

### Lack of HO1 affects the nuclear localization of p65

To better understand what aspect of NF-κB function is influenced, we compared the effects of TNFα stimulation in WT and KO-Hmox1 cells. Western blots (Fig. 4A) showed no differences in total protein levels of TNF receptor-1 (TNFR1) and NF-κB p65 subunit between WT and KO-Hmox1 fibroblasts. Moreover, phosphorylation of the NF-κB inhibitor-α (IκB-α; Ser32/36) and the p65 subunit (Ser536), which is a marker of NF-κB activation, was similarly strong in cells of both genotypes 30 minutes after TNFα treatment (Fig. 4A). Also, immunocytochemical staining (Fig. 4B, C) did not reveal significant differences in the total level of p65 in cells, confirming the results of western blots (Fig. 4A). Finally, using luciferase reporter assay we noticed only subtle differences in functional activation of NF-κB, as the response to TNFα in KO-Hmox1 cells, although visible, did not reach statistical significance (Fig. 4D).

**Figure 4.**
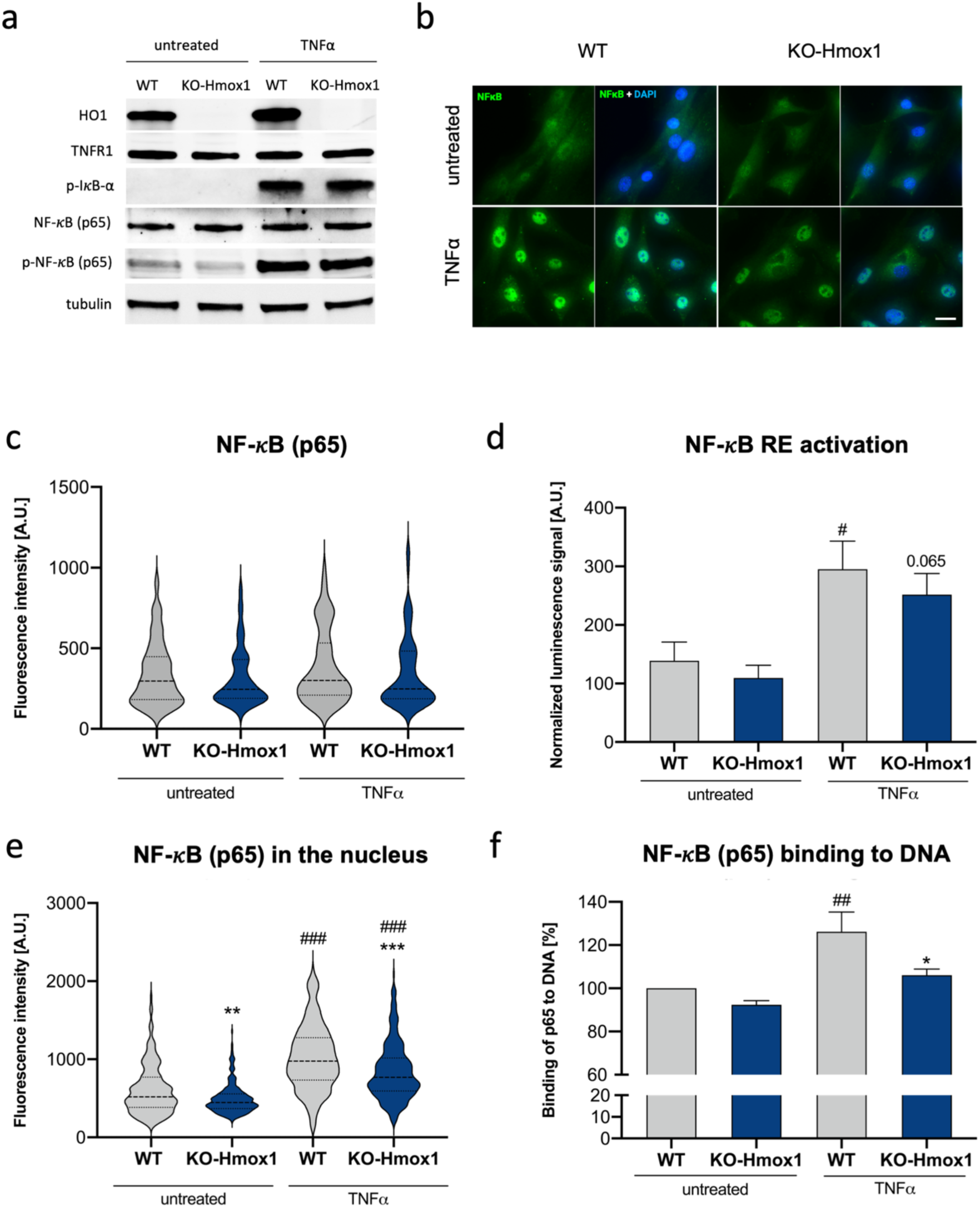
The absence of HO-1 impacts NF-κB signaling pathway. **a)** Western blotting of HO-1, TNFR1, phospho-IκB-α (Ser32/36), NF-κB (p65), phospho-NF-κB (p65 – Ser536) in WT and KO-Hmox1 fibroblasts untreated or treated with TNFα for 30 minutes. Tubulin was used as a loading control. **b)** Representative images of immunofluorescence staining of NF-κB (p65) (green) and nuclei counterstained with DAPI (blue). Scale bar: 20 μm and **c)** statistical analysis of total NF-κB (p65) protein level. Fibroblasts were treated with TNFα for 30 minutes. N=3. Two-way ANOVA. **d)** Transcriptional activity of NF-κB (p65) measured by luciferase reporter system. WT and KO-Hmox1 fibroblasts were treated with 100 ng/mL TNFα for 5 h. N=3. Two-way ANOVA. **e)** Analysis of nuclear levels of NF-κB (p65) in WT and KO-Hmox1 fibroblasts treated with TNFα for 30 minutes. N=3. Two-way ANOVA. **f)** Trans-AM ELISA of nuclear fraction from WT and KO-Hmox1 fibroblasts, untreated or treated with TNFα for 30 minutes. N=3. Two-way ANOVA.

However, analysis of nuclear p65 staining showed a reduced signal in *Hmox1*-deficient cells, both under control conditions and after TNFα stimulation (Fig. 4E). We also compared p65 binding to specific DNA sequences in nuclear fractions isolated from WT and KO-Hmox1 fibroblasts. TransAM ELISA showed that after 30 minutes of stimulation with TNFα, the binding of p65 to its consensus sequences was weaker in KO-Hmox1 cells (Fig. 4F). We suppose that this effect can reflect the reduced levels of nuclear p65 in the absence of HO1. Taken together, our results suggest some impairment of nuclear retention of p65 in *Hmox1*-deficient cells.

### Nuclear localization of p65 is maintained by PARylation in WT but not KO-Hmox1 cells

HO1 has been reported to co-precipitate with PARP1 and modulate both PARP1-mediated PARylation and PARG-mediated dePARylation [36] [37]. In response to proinflammatory signals, PARylation of p65 reduces its interaction with CRM1 exportin, facilitating nuclear accumulation of NF-κB [38]. Therefore, we tested whether PARylation might play a role in the nuclear retention of p65 in WT and KO-Hmox1 fibroblasts.

To do so, we pretreated cells of both genotypes with olaparib (PARP1 and PARP2 inhibitor) for 1 h and then treated them with TNFα for 30 minutes (Fig. 5). We observed that TNFα-induced nuclear accumulation of p65 in WT fibroblasts was completely abolished by olaparib pretreatment (Fig. 5A,B). Surprisingly, in KO-Hmox1 cells, inhibition of PARylation only slightly reduced the level of p65 in the nucleus (Fig. 5A, C). It suggests that PARylation plays a less important role in the regulation of p65 trafficking in *Hmox1*-deficient cells. One can suppose that lack of HO-1 already disturbs the function of PARP1, thus additional inhibition had no significant effect on already lower nuclear p65 in KO-Hmox1 fibroblasts.

**Figure 5.**
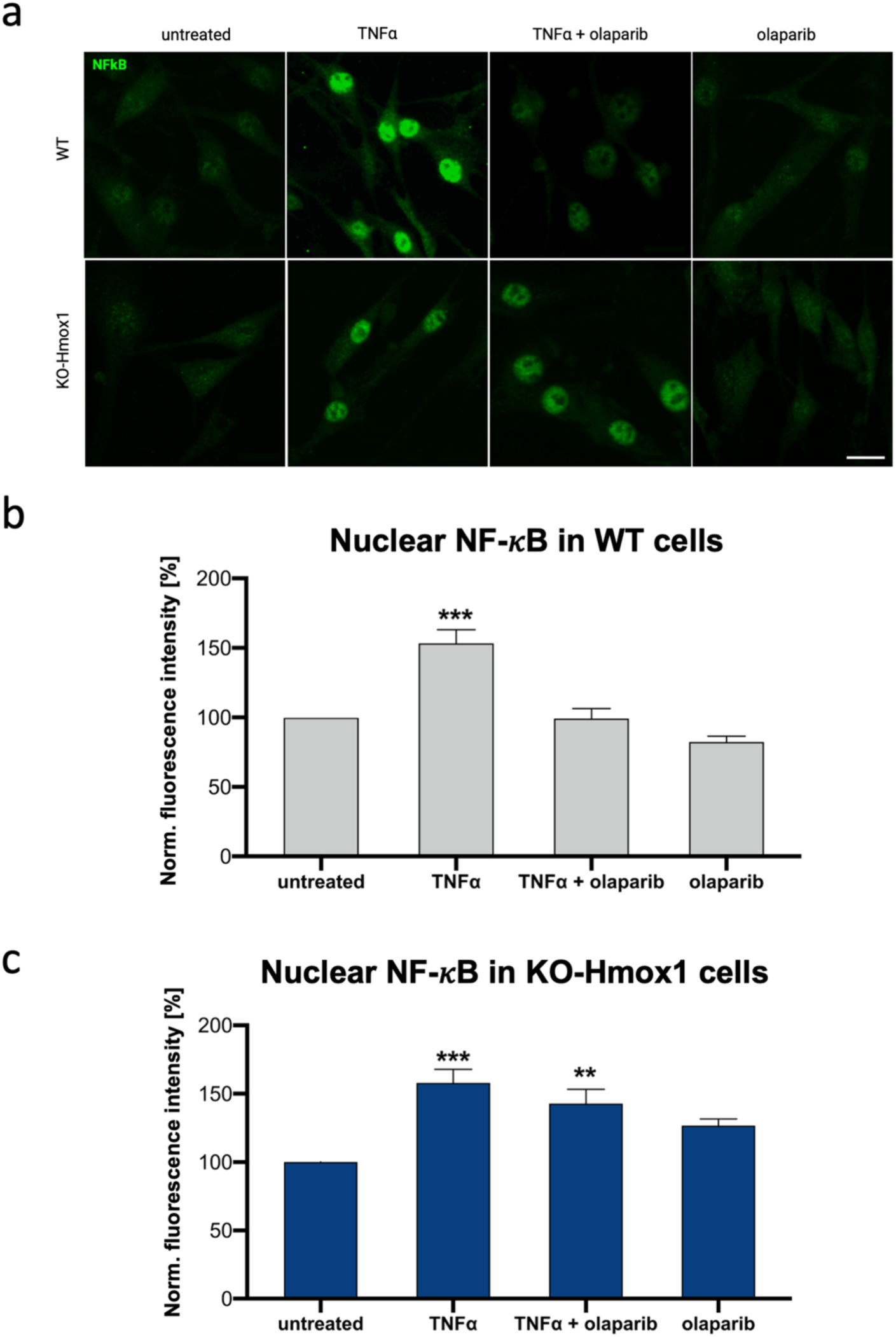
The lack of HO1 affects PARP1-dependent nuclear accumulation of p65. **a)** Representative images of immunofluorescence staining of NF-κB (p65) microscopy (scale bar: 20 µm) and statistical analysis of p65 nuclear levels in **(b)** WT and **(c)** KO-Hmox1 fibroblasts. Cells were treated with TNFα for 30 minutes and 100 nM olaparib for 90 min. N=3, n=17-20. ANOVA.

We also compared the effect of PARylation on CRM1 and p65 interaction in the presence and absence of HO-1. Immunocytochemical staining showed very similar level of CRM1 in the nuclei of WT and KO-Hmox1 cells, although after 24 h stimulation with TNFα we observed some decrease in *Hmox1*-deficient fibroblasts (Fig. 6A). Then, we used proximity ligation assay (PLA) to analyze whether olaparib affects CRM1-p65 colocalization. To avoid the influence of changes in CRM1 expression, we performed PLA in unstimulated cells. In such cells, nuclear localization of p65 was lower in KO-Hmox1 cells (Fig. 4E), despite similar total p65 level (Fig. 4C). The nuclear signal of PLA was also weaker in KO-Hmox1 cells (Fig. 6B). Interestingly, we observed a significant reduction in nuclear colocalization of CRM-p65 in WT cells treated with olaparib, whereas *Hmox1*-deficient cells were insensitive to such treatment (Fig. 6B). This observation confirms that the effects of PARylation are abolished in the absence of HO-1.

**Figure 6.**
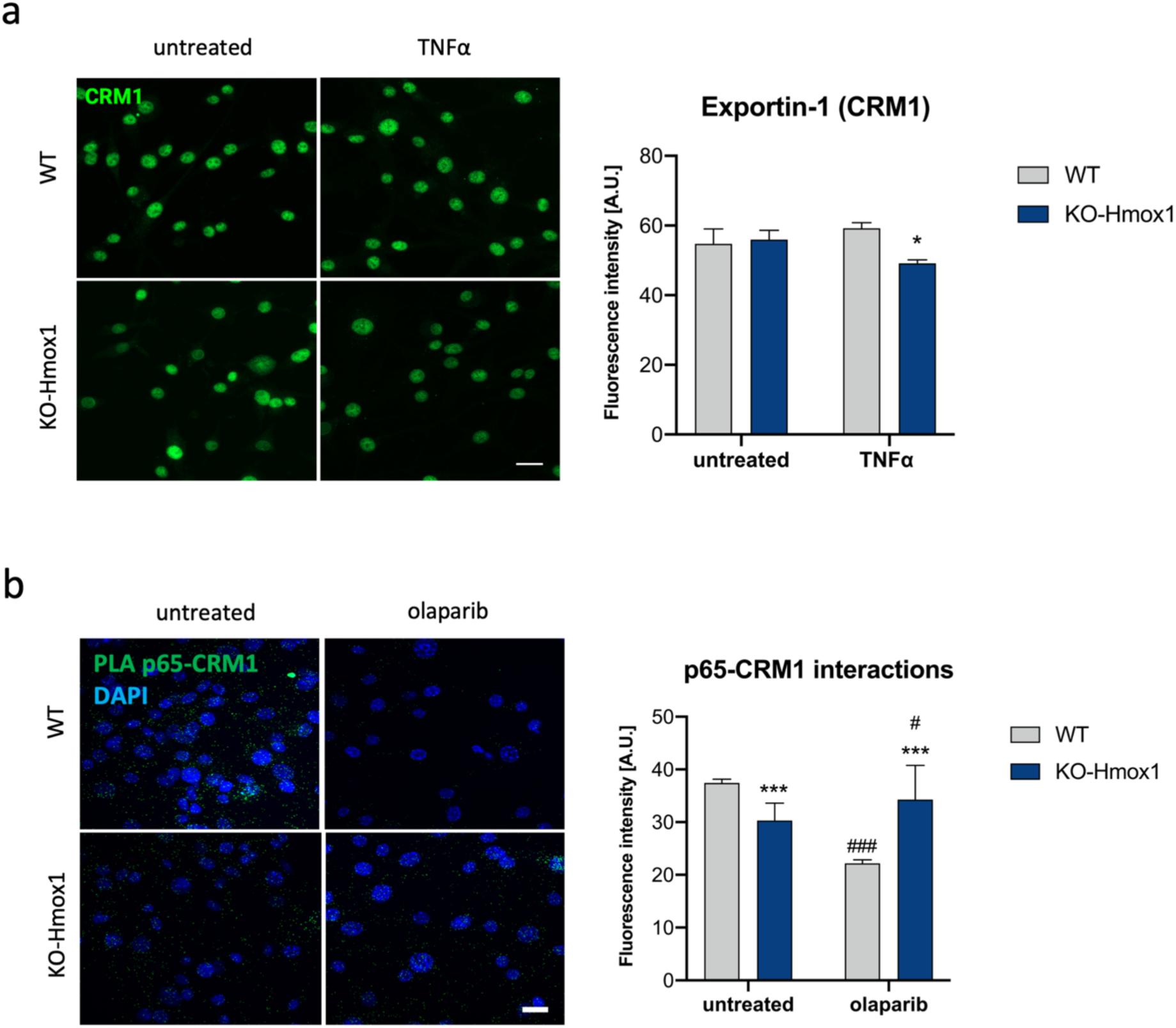
The effect of PARylation on CRM1 and p65 interaction in the cells lacking HO-1. **a)** Representative images of immunofluorescence staining of CRM1 (scale bar: 10 μm) (left) and statistical analysis of its levels in fibroblasts treated with TNFα for 24 h. N=3, two-way ANOVA. **b)** Left panel represents sample images of p65 and CRM1 colocalization (green *foci*) visualized by proximity ligation assay (PLA) in WT and KO-Hmox1 fibroblasts, untreated or treated with olaparib (100 nM) for 24 h. Cell nuclei were counterstained with DAPI (blue). Scale bar: 10 μm. Right panel represents quantitative analysis of fluorescence signal of PLA measured in the nucleus. * - WT vs KO-Hmox1, # - untreated vs treated. N=3, n=20. Two-way ANOVA.

### Reduced nuclear accumulation of STAT1 in KO-Hmox1 cells

To check whether the observed relationship applies only to the NF-κB pathway or is more general, we investigated the effect of *Hmox1* deficiency on STAT1 and STAT2 transcription factors, which are crucial for the interferon response. IRF9, which was upregulated in both WT and KO-Hmox1 cells in response to TNFα (Fig. 3C), forms heterotrimeric ISGF3 complex with STAT1 and STAT2, recognizing ISRE sequences in ISGs and participating in late IFN-I response. STAT1 forms also homodimers, as a downstream effect of INFAR activation [18]. Importantly, STAT1 transcriptional activity is dependent on PARylation [25].

Total levels of STAT1 and STAT2 were similar in cells of both genotypes cultured under control condition. They were also similarly increased after TNFα stimulation (Fig. 7A-C). As with p65, we observed reduced nuclear accumulation of STAT1 in both untreated and TNFα-stimulated KO-Hmox1 cells (Fig. 7C). STAT2 level was not reduced in KO-Hmox1 cells. Conversely, under control conditions, the nuclear staining for STAT2 was even stronger (Fig. 7E). Therefore, in the next step, we focused on STAT1, which shows a similar regulation pattern as p65.

**Figure 7.**
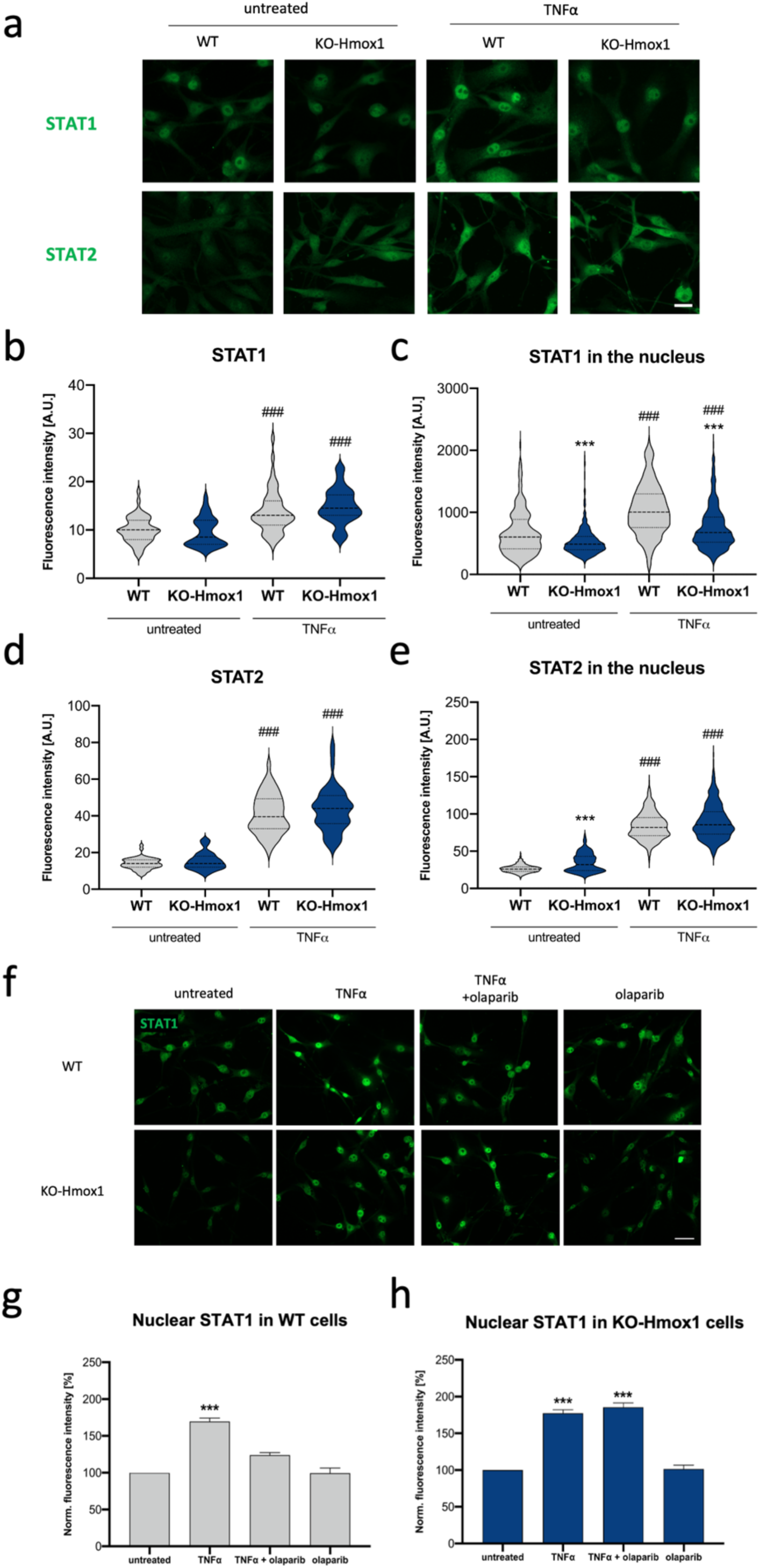
HO1-deficiency results in decreased nuclear accumulation of STAT1. **a)** Representative images of STAT1 and STAT2 immunostaining in WT and KO-Hmox1 fibroblasts in control conditions and after TNFα stimulation. Scale bar: 20 µm. Quantitative analysis of STAT1 levels **b)** in cells and **c)** in the cell nucleus only. Fibroblasts were treated with TNFα for 24 h. N=3. One-way ANOVA. Quantitative analysis of STAT2 levels **d)** in cells and (**e)** in the cell nucleus only. Fibroblasts were treated with TNFα for 24 h. N=3. Two-way ANOVA. **f)** Representative images of immunofluorescence staining of STAT1 (Scale bar: 40 µm) **g)** quantitative analysis of STAT1 nuclear levels in fibroblasts WT and **(h)** KO-Hmox1 treated for 24 h with 10 ng/mL TNFα and 100 nM olaparib (1 h of preincubation) both individually or in combination. N=3, n=28-30. ANOVA.

To investigate the importance of PARylation, similarly to p65 analyses, we preincubated cells of both genotypes with olaparib for 1 h and then stimulated them with TNFα. The obtained results (Fig. 7F-H) were strikingly similar to those for p65 (Fig. 5). Namely, TNFα-induced nuclear accumulation of STAT1 was completely blocked by olaparib pretreatment in WT cells (Fig. 7G), whereas in KO-Hmox1 cells, olaparib showed no effect on nuclear STAT1 (Fig. 7H). Thus, the nuclear accumulation of some transcription factors, such as NF-κB and STAT1 relies on different mechanisms in *Hmox1*-competent and *Hmox1*-deficient cells. In the absence of HO1, cells rely less on PARylation-regulated processes.

### Hmox1-deficiency increases envelope permeability

Our results indicate that *Hmox1* deficiency affects nuclear accumulation of some transcription factors, possibly through several mechanisms, including deregulation of PARylation-dependent pathways. Macromolecular transport between the nucleus and cytoplasm is permitted by the nuclear envelope, not only by the nuclear pore complex but also by structural components, including lamina [39], [40]. There is direct and indirect crosstalk between PARylation and nuclear envelope proteins, with lamin A/C as one of the targets [41], [42]. Therefore, we checked whether *Hmox1* deficiency could also affect lamin A/C levels. Indeed, immunofluorescence staining showed a decreased level of lamin A/C in KO-Hmox1 cells, both under control conditions and after TNFα treatment (Fig. 8A). Given this and the alterations in CRM1 function (Fig. 6B), we expected that Hmox1 deficiency might generally affect nuclear envelope permeability.

**Figure 8.**
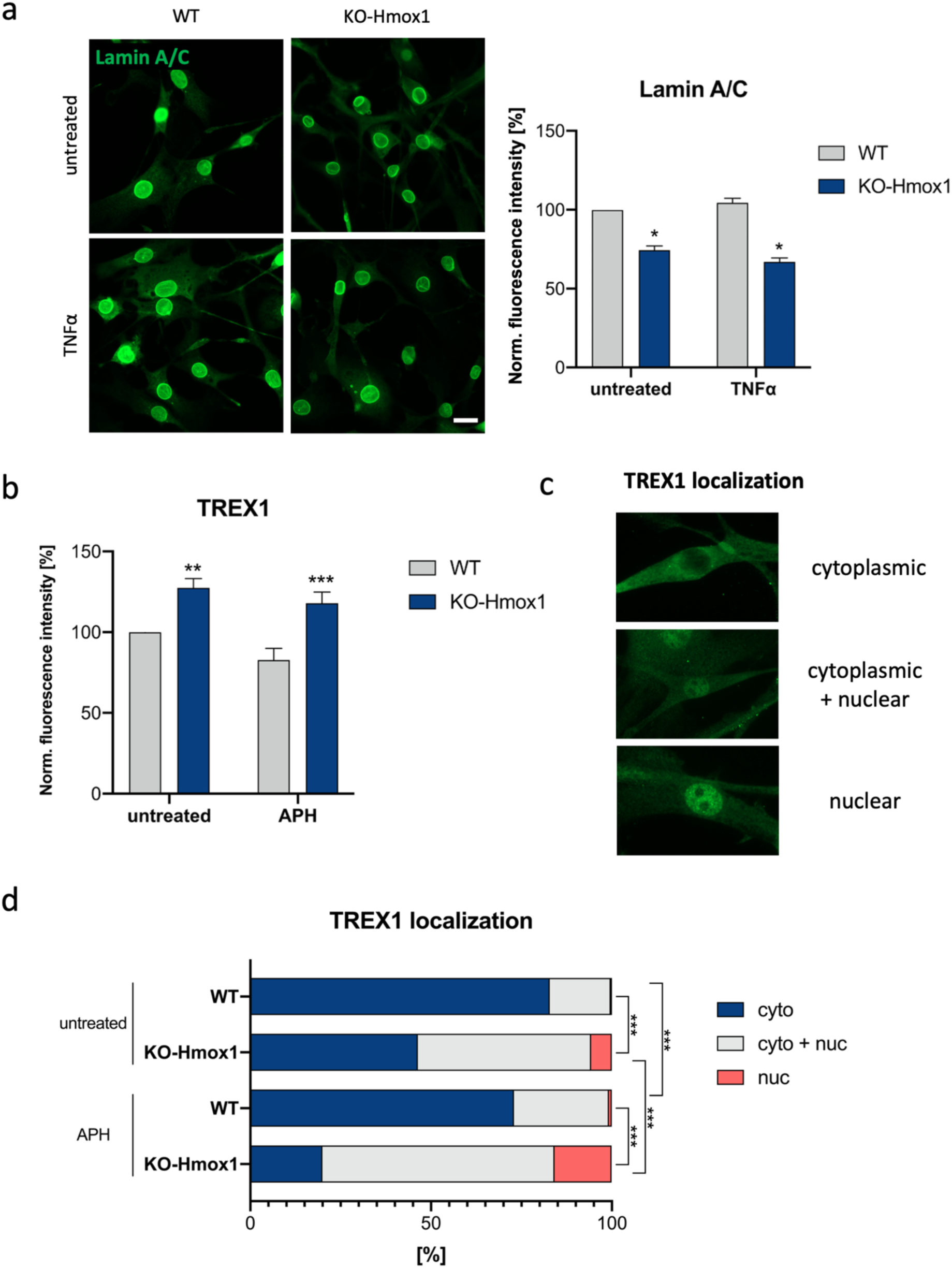
Permeability of nuclear envelope is increased in *Hmox1*-deficient cells. **a)** Representative images (left) and quantitative analysis of immunofluorescence staining of lamin A (right) in fibroblasts treated with TNFα for 24 h. N=3. Two-way ANOVA. Data were normalized to WT untreated cells. **b)** Quantitative analysis of immunofluorescence staining of TREX1 in fibroblasts treated with APH for 24 h. N=3. Two-way ANOVA. **c)** Representative images of TREX1 depicting protein localized in: cytoplasm only (cyto), both in cytoplasm and nucleus (cyto + nuc) and only in the nucleus (nuc). **d)** The percentage of cells expressing TREX1 protein in specific cell compartments of WT and KO-Hmox1 fibroblasts treated with 10 ng/mL APH for 24 h. N=3. Chi^2^ test.

The nuclear localization of TREX1 exonuclease has emerged as an indicator of nuclear envelope permeability, particularly in the context of cellular stress and DNA damage. Under physiological conditions TREX1 is associated with endoplasmic reticulum and its presence in the nucleus may reflect compromised nuclear envelope integrity [43]. We recently demonstrated that a universal effect of *Hmox1* deficiency is replication stress, observed among others in HSCs isolated from the bone marrow of *Hmox1* KO mice and in lymphoblastoid cells from an *HMOX1*-deficient patient [3]. We also observed increased DNA damage response (DDR), indicated by γH2AX staining, in KO-*Hmox1* fibroblasts, both under control conditions and after induction of genotoxic or replication stress using etoposide or aphidicolin, respectively (Fig. S2A).

We performed immunocytochemical staining for TREX1 in WT and KO-Hmox1 cells (Fig. 8B). Total level of TREX1 was increased in unstimulated *Hmox1*-deficient cells when compared to WT counterparts. A similar increase was visible in cells under replication stress, induced with aphidicolin (Fig. 8B). It is worth noting that TREX1 belongs to ISGs and its expression increases in response to the presence of cytosolic DNA [44]. CGAS and STING1, two other proteins of this pathway, were unaffected (Fig. S2B, C). Next, we analyzed the localization of TREX1. For this purpose, we distinguished three possibilities of TREX1 presence (Fig. 8C): i) mainly in the cytoplasm (cyto); ii) in both the cytoplasm and the nucleus (cyto + nuc); iii) mainly in the nucleus (nuc). As expected, in WT fibroblasts TREX1 was visible mostly in the cytoplasm (80% of cells). In response to aphidicolin, the fraction of cyto-nuc cells increased significantly (Fig. 8D), but still the percentage of cells with dominant nuclear localization of TREX1 was negligible. Interestingly, in KO-Hmox1 cells, under control conditions, TREX1 was present in the cytoplasm only in 46% of cells, whereas in 48% of cells it was localized both in the cytoplasm and nucleus, and in 6% mainly in the cell nucleus (Fig. 8D). Moreover, aphidicolin further increased TREX1 cyto + nuc (64%) and nuc (16%) localization. This observation is consistent with the supposition that the lack of HO1 is associated with increased permeability of the nuclear envelope.

## Discussion

In previous studies, we have observed that in infection-free *Hmox1* KO mice, there is an increase in the expression of interferon-stimulated genes (ISGs) in many cell types, despite no increase in interferon levels [8], [9], [45]. Current research leads to conclusion that although cell-autonomous upregulation is possible for some genes (such as *Ifi27*), in most cases there is an extrinsic regulation of ISGs *in vivo*, most likely as a response to pre-inflammatory state and increased production of proinflammatory cytokines characteristic of KO-Hmox1 mice. Concomitantly, however, under inflammatory conditions induced by TNFα exposure *in vitro*, we found that the response of *Hmox1*-deficient cells is weakened. Our results show that HO1 expression improves nuclear envelope integrity, facilitates nuclear retention of NF-κB and STAT1, and maintains sensitivity to PARylation-mediated regulations of nuclear-cytoplasm transport.

HO1 is a heme-degrading enzyme, induced in response to oxidative stress and inflammation, which has been described for years as cytoprotective and anti-inflammatory due to its products and by direct interaction with proinflammatory transcription factors [35]. In mice, *Hmox1* deficiency leads to microcytic and hemolytic anemia, observed already in pups and enhanced in aged animals, as confirmed by very low level of circulating hepcidin [46]. This is mainly caused by disturbed iron metabolism and depletion of CD163^+^ macrophages responsible for removal of senescent erythrocytes, typical of KO-Hmox1 mice [15,16]. Importantly, heme released during hemolytic stress into the extracellular space can bind as a partial agonist to membrane TLR4 to activate both the NF-κB-dependent inflammation and the IRF-dependent IFN-I response [49].

TLR signaling can be classified as either MyD88-dependent which drives the induction of inflammatory cytokines, or TRIF-dependent responsible for the induction of IRF-I response as well as inflammatory cytokines. TLR4 is the only TLR that activates both the MyD88- and TRIF-dependent pathways [33]. It activates IRF3, IRF7, or IRF8, and thereby induces the ISGs [50]. We initially suspected that the increased ISG expressions observed in KO-Hmox1 mice may be the result of hemolytic stress and TLR4 activation by heme.

To examine the effect of hemolytic stress on ISGs, in this study we stimulated WT and KO-Hmox1 fibroblasts with hemoglobin. Additionally, we used 5-aminolevulinic acid, a heme synthesis substrate, to test the influence of intracellular free heme overload. Recently, we showed that the lack of HO1 increases the free heme pool after ALA treatment [3]. Interestingly, we did not observe any differences in ISG expressions in response to Hb or ALA, which indicates that hemolytic stress or increased labile heme probably are not responsible for the upregulation of ISGs observed *in vivo*. Moreover, the induction of DNA damage response or replication stress, both enhanced by the lack of HO1 in HEK293 cells [3] and murine bone marrow-derived HSCs [5], also failed to induce expression of ISGs in mouse primary fibroblasts, regardless of HO1 status.

HO1 reduces oxidative stress by removing the excess of free heme, which may be prooxidant. However, untreated WT and KO-Hmox1 fibroblasts had similar levels of ROS and lipid peroxidation, and additional oxidative stress did not induce ISGs. This resembles the pattern we observed in WT and KO-HMOX1 HEK293 cells [3]. What is more, bone marrow mesenchymal stromal cells isolated from KO-Hmox1 mice were able to quickly activate antioxidant defense when challenged with hemin [51]. This suggests that oxidative stress is not the main cause of IFN-I response. The only stressor inducing ISGs expression (*Irf1, Irf7, Irf9, Ifitm3, Oas1g, Oasl1*) in fibroblasts cultured *in vitro* was TNFα, a proinflammatory cytokine. Therefore, we believe that the induction of ISGs in KO-Hmox1 mice is due to a pre-inflammatory state and increased production of pro-inflammatory cytokines, including TNFα.

The involvement of HO1 in the regulation of interferon response has been reported previously. Namely, it was shown that HO1 is required for TLR4-induced production of IFN-I and expression of IRF3 target genes in macrophages [19]. IRF3 exists in two forms, a monomeric form that is mainly exported to the cytoplasm, and a phosphorylated dimeric form that is sequestered in the nucleus. Interestingly, HO1 was suggested to directly interact with IRF3 leading to its increased phosphorylation and subsequent nuclear retention [19]. Although the underlying pathway is different, the observation of increased sequestration of IRF3 in the nucleus is consistent with our results indicating increased nuclear retention of NF-κB, STAT1 (present study), and p53 [3], which may suggest a more universal link between HO1 and cytoplasm-nuclear exchange.

The modifying effect of HO1 may result from different and not mutually exclusive mechanisms. HO1 was recently shown to positively regulate IFN-I signaling in viral infections, possibly due to increased IRF3 nuclear retention [20]. In contrast, it inhibited IFN-I and ISRE promoter inductions after H_2_O_2_ treatment, due to the promotion of IRF3 degradation via an autophagosome-dependent pathway. This was dependent on HO1 enzymatic activity and iron availability [20]. However, in our experimental setting, we did not observe any effect of H_2_O_2_ on ISG expressions.

It is known that stimulation of human fibroblasts or macrophages with TNFα significantly increases IFN-I response [52] but mutual regulations of IFN-I pathway and inflammation are cell-type dependent [53]. Of note, KO-Hmox1 mice have higher levels of circulating MCP1, TNFα, and several other proinflammatory cytokines, whereas elevated expression of HO1, resulting from *HMOX1* promoter polymorphism, provides better resistance to oxidative stress and reduces the inflammatory response in the human population [54]. This reveals that the final output of HO1 is to mitigate inflammation. However, the regulation of individual components of the inflammatory response may be more complex and cell-type specific.

Furthermore, iron, the end-product of HO-1 enzymatic activity causes the inhibition of NF-κB nuclear translocation in primary prostate cells [55]. Iron was also reported to inhibit the phosphorylation of p65, thereby reducing NF-κB function in endothelial cells [56]. Docking analysis showed that HO-1 might also interact with p65 to decrease its binding to DNA [35]. Finally, no significant effect of HO1 on NF-κB was observed in macrophages [19].

Interestingly, primary mouse KO-Hmox1 fibroblasts cultured *in vitro* demonstrated weaker response to exogenous TNFα stimulation. This observation initially suggested a disturbance in TNF receptor signal transduction in the absence of HO1. However, western blot and immunocytochemical staining did not show significant differences in TNFR and total or phosphorylated p65. The key parameter, which was different between WT and KO-Hmox1 cells was the cellular localization of the p65 subunit of NF-κB, which translocates to the nucleus and activates the expression of INF-I genes in response to TNFα. Analysis of p65 subcellular localization and DNA binding showed impaired p65 accumulation and function in the nuclei of KO-Hmox1 cells. This could potentially result from abnormal nuclear transport, including both import and export pathways or faulty nuclear envelope.

A very important universal regulator of transport between the nucleus and the cytoplasm is PARylation. It was reported that PARP1 and NF-κB form a stable, immunoprecipitable nuclear complex [57], [58] and PARylation of p65 was a critical determinant for its interaction with nuclear export protein CRM1, promoting nuclear retention of NF-κB [59]. The mechanism may not be universal, as direct interaction with p65 was not observed in *Trypanosoma cruzi*-infected cardiomyocytes, where PARP1-mediated PARylation of p65-interacting nuclear proteins promoted NF-κB activation and cytokine gene expressions [58]. Importantly, inhibition of PARylation reduced the nuclear localization of p65 in such cells [58] and in smooth muscle cells after TLR4 stimulation [60], an effect very reminiscent of what we observed in wild-type fibroblasts treated with olaparib.

Moreover, we found a very similar pattern as for p65 also for STAT1. Namely, HO1 enhanced nuclear localization of STAT1 in primary mouse fibroblasts, which is consistent with a previous report showing higher STAT1 activity in *Hmox1*-expressing cells [6]. Similar to p65, inhibition of PARylation abolished the TNFα-induced nuclear retention of STAT1. This indicates that the regulatory role of PARylation is not limited to a specific transcription factor but is more general.

Indeed, PARP-1 has been implicated in controlling the subcellular localization of NF-κB, STAT1, and p53 [60], [61]. There is no known relationship between PARP1 and proteins of the importin system, such as importin a3 and a4 [60]. In contrast, PARP1 expression and activity affect the cytosolic levels of CRM1 and dynamics of CRM1 trafficking, which is required for the nuclear export of NF-κB, p53, and STAT1 [60]. PARylation of p53 or p65 NF-κB by PARP1 reduces their interaction with CRM1, hence rendering the transcription factors resistant to export and promoting their retention in the nucleus [38], [60], [61]. Of note, we observed that olaparib strongly affects colocalization of p65 and CRM1 in wild-type fibroblasts, although the functional effects of these changes measured 24 hours after olaparib administration cannot be unequivocally interpreted. Unexpectedly, the effects of olaparib on the TNFα-induced p65 and STAT1 nuclear retention as well as p65-CRM1 colocalization are totally abolished in the absence of HO1.

Thus, a model has emerged in which PARylation of stress-responsive transcription factors blocks their nuclear export, thereby promoting their accumulation in the nucleus [38]. Our results indicate that these regulatory pathways do not function properly in *Hmox1*-deficient cells.

For now, we can only speculate about the reasons for this phenomenon. Interestingly, HO1 can co-precipitate with PARP1, suggesting a direct protein-protein interaction [37]. Based on docking modeling and pull-down experiments, it was proposed that HO1 binds to the regulatory helical domain (HD) of PARP1 [37], which might result in the opening of its folded structure, leaving PARP1 in a continuous activation state [37]. Additionally, HO1 was suggested to bind to the PARG protein, reducing its dePARylating activity [62]. These assumptions indicate that in the absence of HO1, the PARylation level may be lower, which could consequently facilitate the export of transcription factors from the nucleus.

However, our recent studies did not detect such a relationship. Proximity ligation assay confirmed a colocalization of PARP1 and HO1, but PARP1 accumulated at the sites of DNA damage more rapidly in the absence of HO1 [3]. Furthermore, PAR formation at the site of laser micro-irradiation was also more rapid in HO1-deficient cells. On the other hand, autoPARylation of purified PARP1 protein was comparable in the presence or absence of HO1. We concluded that the effect of HO1 deficiency on PARylation was rather associated with PARP1 cellular motility, but not with a substrate availability or direct protein-protein interaction modulating PARP1 enzymatic activity [3]. We also demonstrated that the effects of HO-1 on p53 are more related to the regulation of free heme than to PARylation [3].

Another putative trigger of IFN-I response and ISG expressions can be a pool of cytosolic DNA, produced intrinsically during cell proliferation and DNA repair. Their lengths range between 100 and 1000 nt for double-stranded DNA (dsDNA) and IFN-I pathway is induced by fragments longer that 24 nt, irrespective of sequence [63], [64]. It appears that the main intrinsic source of cytoplasmic DNA is replication stress and fork retention that activate DNA damage response (DDR). Recently, we demonstrated that HO1 co-localizes with DNA G4 structures both in the nucleus and cytoplasm and plays a role in the regulation of G4 unwinding [5]. Moreover, lack of *Hmox1* in proliferating cells is associated with enhanced formation of DNA G4 structures and replication fork stalling [3]. Here, in KO-Hmox1 fibroblasts we also saw an increased level of DDR under control conditions and after stimulation with etoposide (genotoxic stress) and aphidicolin (replication stress). Unfortunately, we were not able to reliably compare cytoplasmic DNA between WT and KO-Hmox1 fibroblasts. However, experiments in murine iPS cells reveal that the lack of HO1 leads to increased levels of cytoplasmic DNA after aphidicolin stimulation [data not shown].

Finally, abnormal nuclear localization of crucial transcription factors as well as cytoplasmic leakage of DNA might be caused by deregulated permeability of nuclear envelope. The inner membrane of nuclear envelope is stabilized by lamins, whereas the outer membrane constitutes a continuum with the endoplasmic reticulum [65], the primary cellular localization of HO1. In laminopathies, disintegration of pores in the interphase leads to disturbances in proteins localization [66]. Our data show that *Hmox1*-deficient cells have lower levels of lamin A, a key protein for maintaining nuclear structure and mechanical stability.

A marker of impaired envelope integrity may be the presence of TREX1 exonuclease in the nucleus [43], where it cuts chromatin leading to DNA damage [67]. In *Hmox1*-deficient cells the level of TREX1 is increased, what may contribute to elevated dsDNA breaks.

Importantly, KO-Hmox1 fibroblasts also had higher nuclear localization of TREX1, suggesting disturbances in the integrity of the nuclear envelope and potentially additional DNA cutting. Other proteins that cooperate with TREX1 in the type-I interferon response are STING (stimulator of interferon genes) and CGAS (cyclic GMP-AMP synthase) [68]. STING is a protein of the inner nuclear membrane stabilized, among others, by lamin A [69]. It was reported that STING can translocate to the nucleus and, additionally, co-precipitate with PARP1. CGAS in the nucleus is largely bound to chromosomes [51]. However, we did not notice any significant differences in levels and location of STING and CGAS between WT and KO-Hmox1 fibroblasts.

## Summary

In conclusion, our results indicate that the increased expression of ISGs we previously observed in *Hmox1*-deficient mice is rather induced extrinsically at the organismal level and may be mediated by increased production of proinflammatory cytokines such as TNFα. In cells, *Hmox1* deficiency leads intrinsically to an attenuated response to TNFα and impaired nuclear retention of NF-kB and STAT1. *Hmox1*-deficient cells are less dependent on PARylation as a regulatory mechanism for transcription factor trafficking and are more susceptible to disruption of the nuclear envelope integrity.

## Author contributions

Conception and design: AJ, PC, WN; Acquisition of data: PC, KB, ECH, JW, AS, GS; Analysis and interpretation of data: PC, AJ, KB, ECH, WK, MZ, KS, AS, WN; Writing, review, and revision of the manuscript: PC, AJ, WN, WK, MZ, AS.

## Acknowledgments

We thank Dr. Katarzyna Miękus (Department of General Biochemistry Faculty of Biochemistry, Biophysics and Biotechnology Jagiellonian University) for providing Leica DMI6000 fluorescence microscope. We thank Malopolska Centre of Biotechnology for providing Zeiss LSM 880 confocal microscope and Dr. Paweł Hermanowicz for assist.

## Data availability

Data will be made available on request.

