## Supplementary material for "Effect of heme oxygenase-1 on the expression of interferon-stimulated genes": Supp. Fig. 1

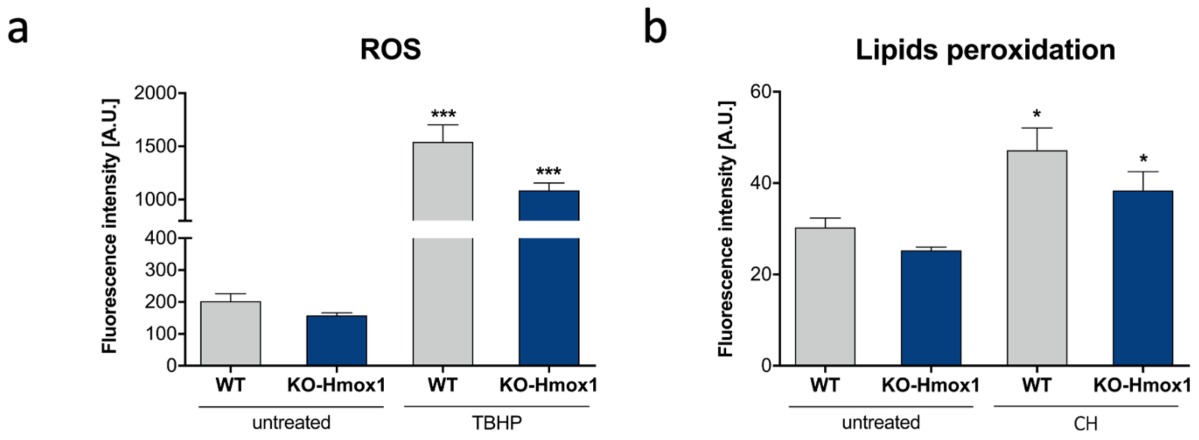


**Fig. S1 a)** ROS levels in WT and KO-Hmox1 fibroblasts. TBHP (200 μM, 30 minutes) was used as a positive control. N=3. Two-way ANOVA. **b)** Lipids peroxidation in WT and KO-Hmox1 fibroblasts. Cumene hydroperoxide (CH, 100 μM, 24 h) was used as a positive control. N=3. Two-way ANOVA.
