## Supplementary material for "Effect of heme oxygenase-1 on the expression of interferon-stimulated genes": Supp. Fig. 2

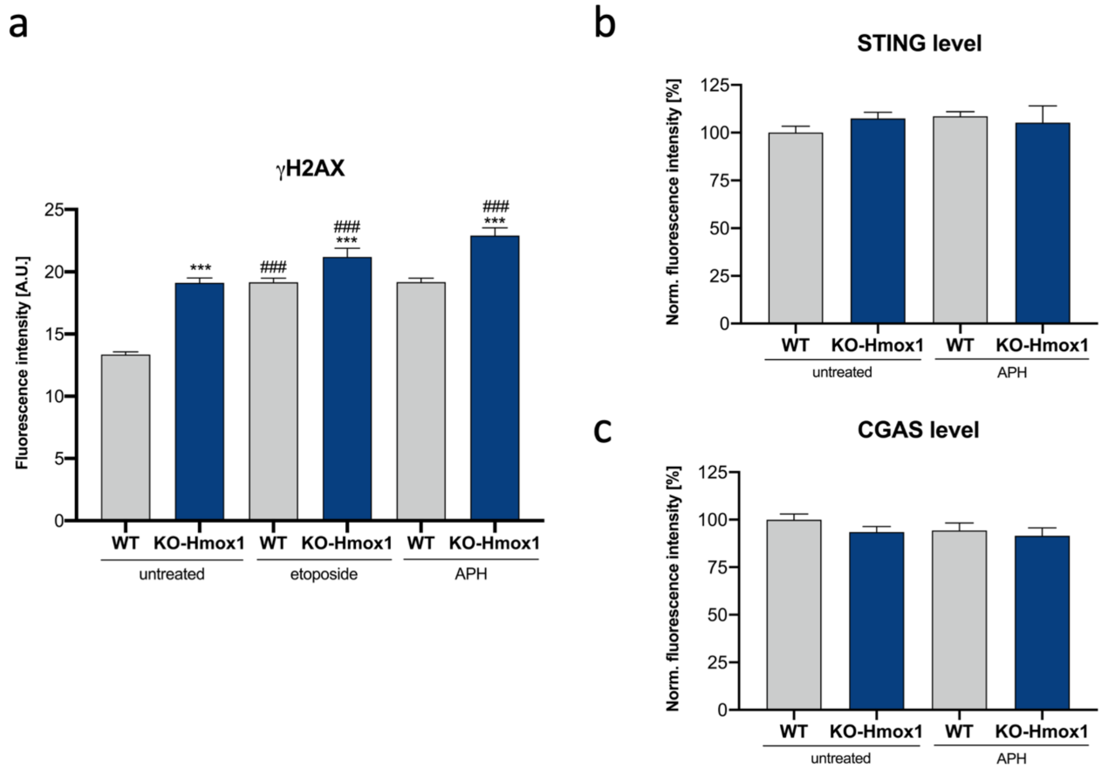


**Fig. S2 a)** Quantitative analysis of γH2AX fluorescence signal in fibroblasts treated with 0.25 μM etoposide or 0.01 μg/ml aphidicolin for 24h. N=3, n=326-792. Two-way ANOVA. * - WT vs KO-Hmox1, # - untreated vs treated. **b)** Quantitative analysis of of immunofluorescence staining of STING and **(c)** CGAS in fibroblasts treated with 0.01 μg/ml aphidicolin (APH) for 24h. N=3. Two-way ANOVA.
